# A chromosome-level, haplotype-resolved genome assembly for the barn owl, *Tyto alba*

**DOI:** 10.64898/2026.03.20.713190

**Authors:** Hugo Corval, Anne-Lyse Ducrest, Marianne Bachmann Salvy, Allison Burns, Alexandros Topaloudis, Céline Simon, Elisa Cora, Daniel Cavaleri, Bettina Almasi, Alexandre Roulin, Christian Iseli, Nicolas Guex, Tristan Cumer, Jérôme Goudet

**Author notes:** corresponding authors. Contact details: Department of Ecology and Evolution, Biophore Building, University of Lausanne, CH-1015 Lausanne, Switzerland. co-senior authors (Nicolas Guex, Tristan Cumer, Jérôme Goudet).

## Abstract

Recent advances in long-read sequencing have enabled near telomere-to-telomere (T2T) assemblies across diverse taxa. However, avian genomes remain challenging due to numerous microchromosomes, small, typically < 20Mb, elements that are gene-, GC-, and repeat-rich. As a consequence, microchromosomes are often missing from genome assemblies. Here, we present a chromosome-level, haplotype-resolved genome assembly for the Western barn owl (*Tyto alba*). Using a trio-binning strategy with Illumina parental reads combined with PacBio HiFi and Oxford Nanopore Technologies data, we generated two phased contig sets. These were scaffolded into 40 linkage groups using a linkage map. Comparative analyses identified unplaced HiFi scaffolds corresponding to microchromosomes, which we integrated into six additional microchromosomes using long reads information. The two assemblies present 46 chromosomes, matching the karyotype of the species. They exhibit strong synteny between parental haplotypes, except for a ∼38 Mb complex region on chromosome 7 containing nested inversions. This high-quality reference provides the first haplotype-resolved and chromosome-level genome for *Strigiformes*, enabling fine-scale studies of structural variation and avian genome evolution.

## Background

In recent years, genomics has entered the telomere-to-telomere (T2T) era. This shift began with the first complete human genome (T2T-CHM13) [1] and quickly expanded to other species, from crops [2,3] to livestock [4] and primates [5]. These milestones were achieved thanks to the advent of long-read sequencing technologies, mostly led by Pacific Biosciences (PacBio) and Oxford Nanopore Technologies (ONT), combined with chromosome conformation contact Hi-C sequencing [6]. With this resolution, assemblies now expose complex regions, such as the major histocompatibility complex (MHC) or the Killer-cell immunoglobulin receptors (KIRs) locus, that were previously poorly characterised due to extreme polymorphism and repetitive structure [7–11].

Although these recent advances have transformed genome assembly, significant challenges remain. One difficulty of genome assembly is reconstructing both parental haplotypes in diploid species rather than a mosaic consensus. Common strategies include trio binning with parental short reads [12,13], Hi-C phasing coupled to long, accurate reads [14,15], and Strand-seq for chromosome length phasing [16,17]. When successful, haplotype-resolved assemblies can improve the discovery of complex structural variants [12,18–20]. Still, phasing can be limited by repeat- and GC-rich regions that promote switches or mis-joins [21], by genomes with large size, high heterozygosity or polyploidy (common in plants) [22], and by practical constraints such as the need for parental samples (trio binning) and the cost/complexity of specialised libraries. Consequently, fully phased references are still less common than haploid assembly in many taxa [11,22].

The genome reconstruction also varies widely across taxa. In vertebrates, bird genomes are difficult to assemble due to the presence of numerous microchromosomes [23,24]. These microchromosomes share features that hamper their reconstruction: they are small, gene-rich, GC-rich and contain a high proportion of repetitive elements [24,25]. Because of these properties, many avian genome assemblies contain fewer chromosomes than the actual karyotype, with microchromosomes frequently missing [24]. Recently, the introduction of Oxford Nanopore ultra-long reads has improved their recovery, leading to more complete assemblies in species such as chicken [26], bustard [27], and mallard [25,28]. Still, adoption of these approaches has been uneven across birds and achieving full and accurate resolution of avian microchromosomes remains a major challenge.

Among owls (*Strigiformes)*, genome assembly poses extra difficulties. Their karyotypes feature many microchromosomes and rearrangements [29], making synteny with well-assembled model species less informative [30]. The Common barn owl (*Tyto alba*) illustrates this difficulty. The species has 46 pairs of chromosomes [31] and a complete genome assembly has yet to be achieved. The current assembly generated using PacBio long reads and Bionano optical mapping consists of 70 “pseudo-haplotypes” where phase information is not conserved, (N50 = 36.03 Mb) [32]. This was a major improvement from the previous version, which contained 21,509 contigs with a N50 of 4.62 Mb [33]. Nevertheless, additional work is needed to reach a chromosome-level assembly that matches the 46 pairs observed in the barn owl karyotype.

In this study, we aim to produce a chromosome-level, haplotype-resolved reference genome for the barn owl (*Tyto alba*). We achieve accurate chromosome phasing through trio-binning using Illumina parental reads. We construct unitigs from HiFi PacBio and ONT data, which we then scaffold into chromosomes using linkage map information [34]. Finally, we leverage the gene-rich nature of microchromosomes to manually curate unplaced unitigs, based on the presence of genes typically found on microchromosomes in other species. With this integrated approach, we deliver a high-quality genomic resource, add a chromosome-level reference of an owl, and resolve previously uncharacterised complex structures.

## Results

### Chromosome-scale, haplotype resolved assemblies

Using PacBio HiFi reads (150.8 Gb, ∼ 119x of paternal and ∼117x of maternal expected coverage, Supplementary Table 1), Oxford Nanopore reads (40.2 Gb, ∼32x of paternal and ∼31x of maternal expected coverage, Supplementary Table 1), and linkage information, we generated two chromosome-level, haplotype-resolved assemblies of the barn owl (*Tyto alba*). Each haplotype comprised 46 chromosomes, matching the expected karyotype (Figure 1; Table 1; Supplementary Table 2). The paternal assembly spanned 1.27 Gbp (N50 = 36.7 Mb; N90 = 15.5 Mb, Figure 1A), and the maternal assembly spanned 1.29 Gbp (N50 = 38.3 Mb; N90 = 15.1 Mb, Figure 1B). Both haplomes showed a slight increase in size compared to previous assembly (GCF_018691265.1: 1.25 Gbp, Table 1). Hi-C contact maps showed strong, chromosome-scale diagonals with no spurious trans-chromosomal signals, consistesnt with high contiguity and correct scaffolding (Figure 1C-D).

**Figure 1.**
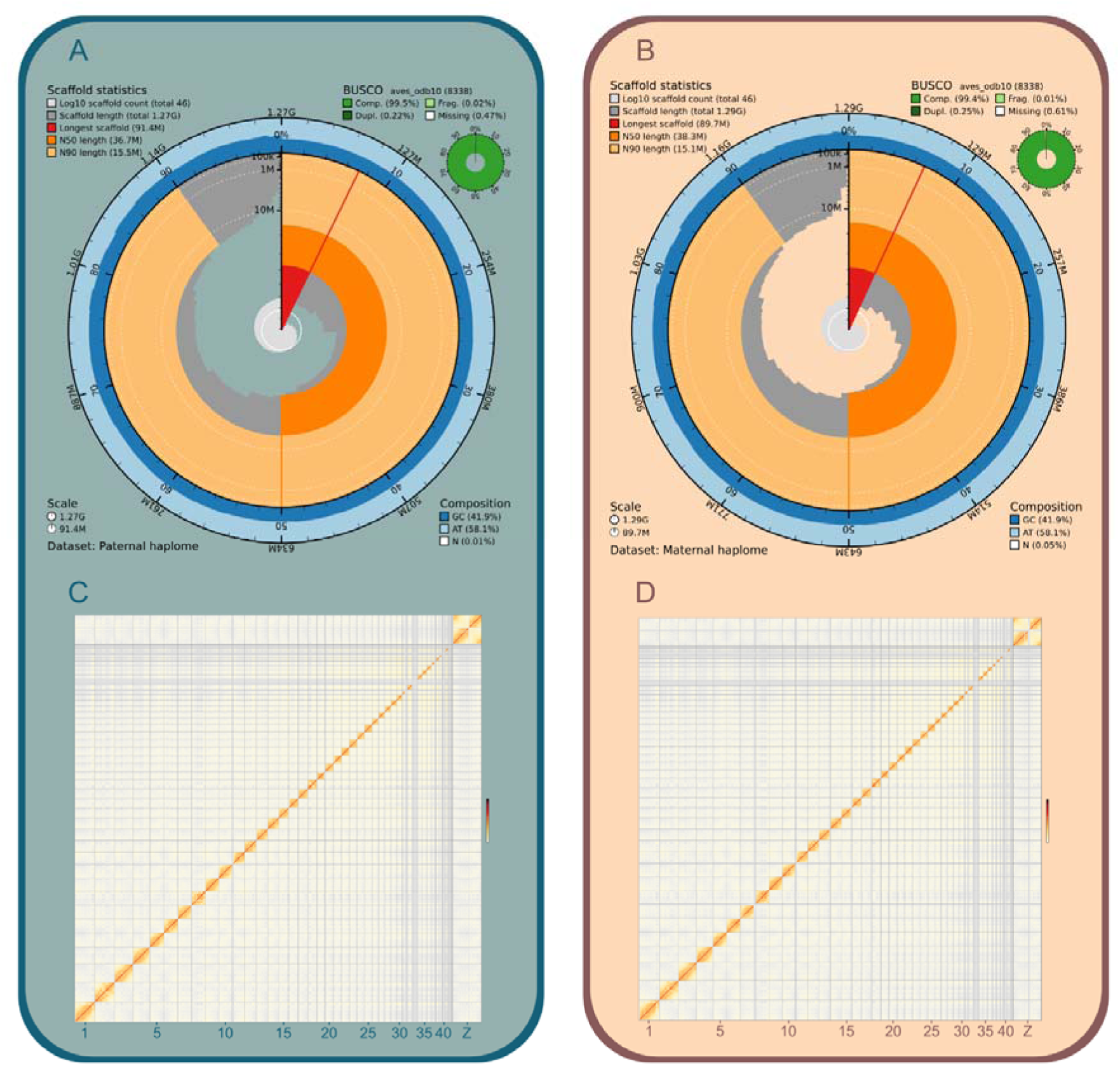
Contiguity, completeness, and chromatin interaction landscape of the paternal and maternal haplotype assemblies. (A-B) Snail plots display the overall structure and composition of the (A) paternal and (B) maternal assemblies. Each circumferential axis is divided into 1,000 bins, each representing 0.1% of the corresponding assembly. Scaffold length distribution is shown in dark grey, with the plot radius scaled to the longest scaffold (highlighted in red). N50 and N90 scaffold length thresholds are indicated in orange and pale orange, respectively. Nucleotide composition is displayed as dark blue for GC, light blue for AT, and white for ambiguous bases (N). A BUSCO completeness summary (Complete, Fragmented, Duplicated, Missing) is provided in the upper-right corner of each panel. (C-D) HiC contact maps for the (C) paternal and (D) maternal assemblies illustrate the genome-wide pattern of spatial interactions. Chromosomes are arranged by size from left to right and from top to bottom, with the Z chromosome positioned last. The red main diagonal represents high-frequency intra-chromosomal contacts, while off-diagonal signals correspond to inter-chromosomal interactions. Interaction frequencies are displayed on a logarithmic scale. Heatmaps were generated using HiContacts.

**Table 1.** Comparison of general assembly statistics across barn owl (Tyto alba) references. Genomes compared include genome_assembly_l500 from Ducrest et al. (2020) and T.alba_DEE_v4.0 from Machado et al. (2021). These reference genomes are shown alongside the two parental haplomes assembled here. Haplomes are each shown twice: a hifiasm column for the unscaffolded contigs, and a final column for the chromosome-scale assembly after filtering, scaffolding and curation. Metrics include total assembly size, counts of contigs, N50/N90, percentage of GC, total Ns and BUSCO completeness.

|  | genome_as<br>sembly_I500 | T.alba_DEE<br>_v4.0 | Paternal |  | Maternal |  |
| --- | --- | --- | --- | --- | --- | --- |
|  | GCF_902150<br>015.1 | GCF_018691<br>265.1 | Hifiasm | Final | Hifiasm | Final |
| **Assembly**<br>**size** | 1,219,191,878 | 1,249,867,532 | 1,317,996,626 | 1,267,676,513 | 1,313,309,923 | 1,285,637,069 |
| **# contigs** | 21509 | 70 | 283 | 46 | 325 | 46 |
| **N50** | 4,615,526 | 36,032,128 | 26,240,345 | 36,723,072 | 24,830,383 | 38,271,680 |
| **N90** | 556,444 | 15,349,174 | 5,753,860 | 15,527,340 | 4,456,682 | 15,107,001 |
| **% GC** | 41.5 | 41.5 | 41.9 | 41.9 | 41.9 | 41.9 |
| **Bp of Ns** | 9,406,517 | 51,418,967 | 0 | 110,000 | 0 | 600,000 |
| **% BUSCO** | 98.4 | 98.5 | - | 99.5 | - | 99.4 |

We quantified phasing accuracy by leveraging k□mers unique to each parental read set. For each haplotype assembly, we computed the switch error rate as the proportion of parental□specific k□mers incorrectly assigned to the opposite haplotype relative to the total number of parental□specific k□mers detected across both assemblies. This analysis indicated that 7.8% of paternal□specific k□mers were misplaced into the maternal haplotype, while 7.5% of maternal□specific k□mers were incorrectly assigned to the paternal haplotype.

As the assembly was constructed from a homogametic male (ZZ), identification of the sexual chromosome could not be based on identification of an heterochromatide unique in each assembly. Instead, to identify the Z chromosome, we used a protein-based homology with other avian species. Species such as chicken (*Gallus gallus*) and zebra finch (*Taeniopygia guttata*) showed that proteins encoded on their Z chromosome mapped predominantly to our largest assembled chromosome (Supplementary Figure 1 and 2). This largest chromosome was also the only metacentric chromosome in the assembly, while all remaining chromosomes were acrocentric (Figure 2B).

**Figure 2.**
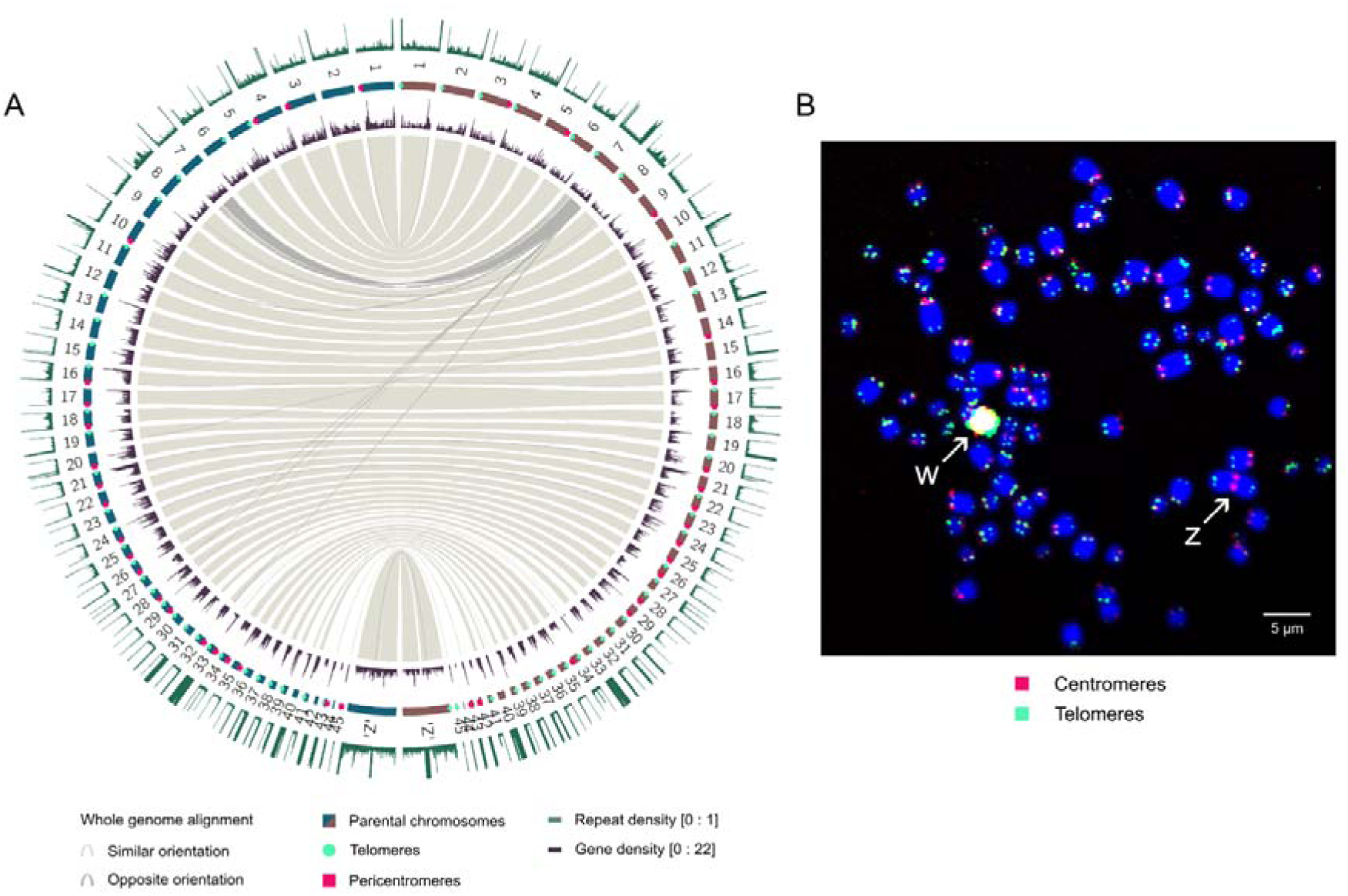
Overview of the paternal and maternal haplomes and their genome features. (A) The Circos plot provides a comparative summary of the genomic characteristics of the paternal haplome (left, dark blue) and maternal haplome (right, burgundy). From the outer to the inner track, the plots show the repeat proportion calculated in 100-kbp windows; the 46 chromosomes of each haplome, with known telomeric and pericentromeric sequences indicated by green circles and red squares, respectively; the distribution of annotated genes, summarised in 100-kbp windows; and the whole-genome alignment between the two haplotypes. Because repetitive elements strongly affect alignment, both haplomes were hard-masked with RepeatMasker prior to alignment. Regions aligning in the same orientation are shown in light grey, and regions aligning in opposite orientation are shown in dark grey. (B) Fluorescence in situ hybridisation (FISH) visualisation of the Common barn owl karyotype of a female individual. Chromosomes are counterstained with DAPI (blue). Centromeric regions are highlighted in red and telomeric regions in green using specific DNA probes (see Material and methods section for details). Sexual chromosomes Z and W are indicated by white arrows.

Relative to prior references, gaps were greatly reduced: 110,000 Ns (0.01%) in the paternal and 600,000 Ns (0.05%) in the maternal haplotype, compared with 51,418,967 Ns (4.11%) in T.alba_DEE_v4.0 (GCF_018691265.1, Table 1). GC content was similar across haplotypes (41.9% each) and only slightly higher than in earlier assemblies (0.1% of increase compared to T.alba_DEE_v4.0). GC profiles were broadly homogeneous between chromosomes on average with a modest increase toward smaller chromosomes (Figure 1A, Figure 1B, Table 1, Supplementary Table 2).

BUSCO assessments indicated high completeness for both haplotypes (paternal: C = 99.5%, F = 0.02%, D = 0.22%, M = 0.47%; maternal: C = 99.4%, F = 0.01%, D = 0.25%, M = 0.61%), representing ∼1% improvement over T.alba_DEE_v4.0 (C = 98.5%).

To evaluate chromosome continuity, we quantified terminal features. In the paternal haplotype, 31 chromosomes contained telomeric sequence in 5p and 21 contained pericentromeric sequence. In the maternal haplotype, 34 and 18 chromosomes contained telomeric and pericentromeric sequence, respectively. Across haplotypes, 24 chromosomes were resolved from telomere to pericentromere, 18 contained only telomeric sequence, 2 centromeric sequence only, and 2 contained neither across haplomes (Figure 2A, Supplementary Table 3).

### Repeat and gene annotation

Custom repeat annotation revealed that repeats constituted 13.68% (paternal) and 14.83% (maternal) of assembled sequence. LINEs were the predominant class (47.81% paternal and 47.07% maternal of total repeats), followed by LTR elements (32.23% paternal and 32.45% maternal, Supplementary Figure 3). Repeats were enriched toward chromosomal ends and in the six smaller chromosomes. On the metacentric Z chromosome, repeat density peaked around the centromere (Figure 2A). The unassembled W chromosome shows intense FISH signals for both telomeric and centromeric probes, indicating a marked enrichment of these repeats (Figure 2B).

Using the NCBI EGAPx workflow with multi-tissue RNA-seq support [33], we annotated 19,720 genes in the paternal haplotype and 20,039 in the maternal haplotype. These haplotype annotations were compared to the previous reference genome (T.alba_DEE_v4.0 - GCF_018691265.1) by mapping back Coding DNA Sequences (CDS) to quantify newly annotated CDS. Newly annotated CDS totalled 908 in the paternal haplotype and 948 in the maternal haplotype and were distributed over the genome, with an enrichment towards the smallest chromosomes (Figure 3D, Figure 3E).

**Figure 3.**
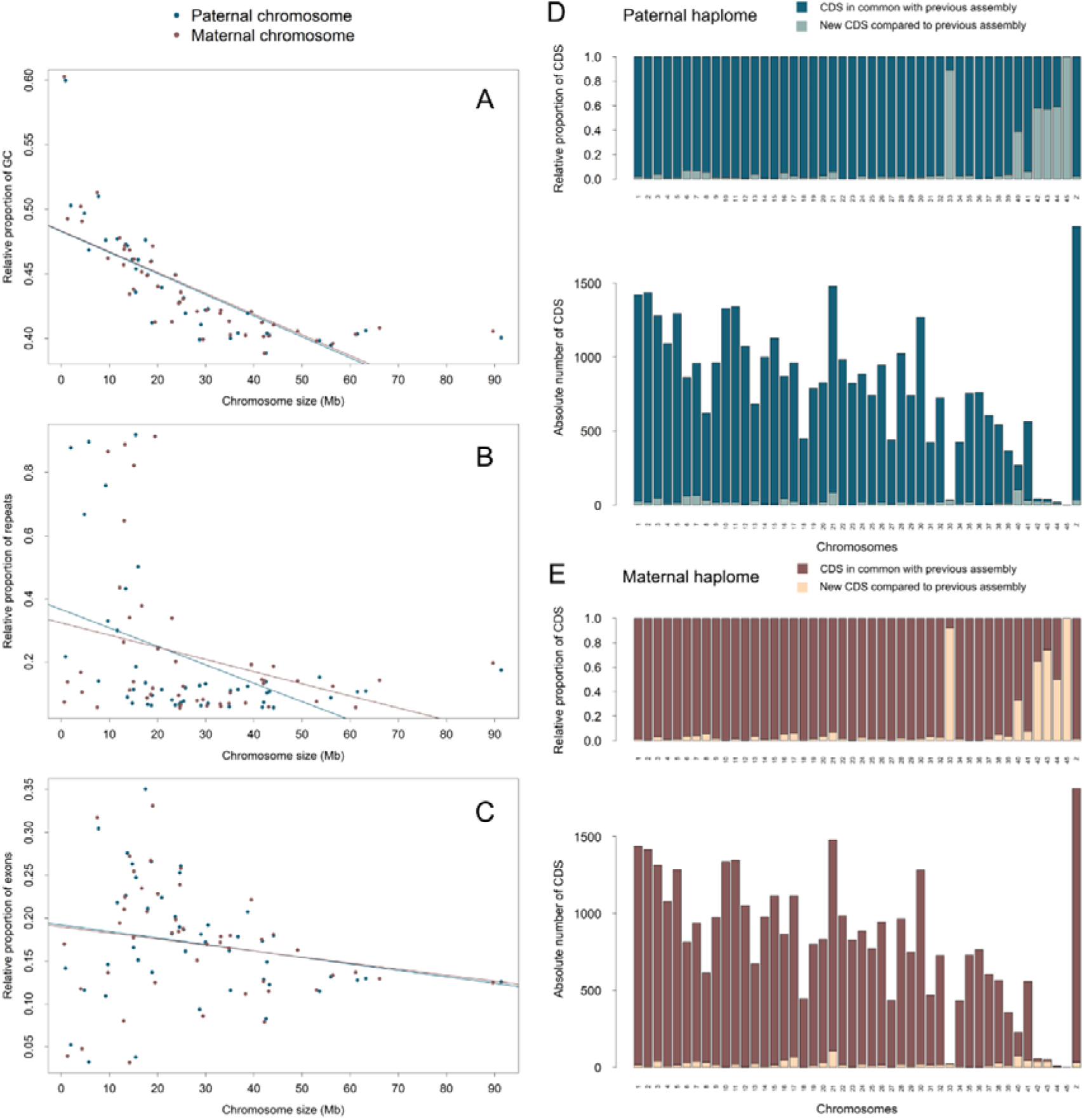
Dot chromosomes and their genomic characteristics. (A-C) Chromosome-wide distributions of GC content (A), repetitive elements (B), and exonic sequence (C) are shown for the paternal (dark blue) and the maternal haplome (burgundy). (D-E) Bar plots illustrate the absolute (lower panels) and relative proportion (upper panels) of newly annotated genes on each chromosome of the paternal (D) and maternal (E) haplomes. Newly annotated genes compared with the previous assembly are depicted in lighter colours.

### Recovery and comparison of microchromosomes

To recover microchromosomes that were missing from the long-reads-based unitigs, we used MicroFinder [24]. This strategy allowed us to re-assign unplaced long-reads-based unitigs to specific microchromosomes and integrate them into the final haplotype assemblies.

We screened 116 unplaced unitigs in the paternal assembly and 119 in the maternal assembly. Based on conserved gene content, we identified 14 paternal and 18 maternal candidates using MicroFinder. We further assembled them in 6 new microchromosomes, that were absent from previous versions (RefSeq: GCF_902150015.1 and GCF_018691265.1).

In contrast to chromosomes already present in the previous assembly, the six newly recovered chromosomes are characterized by distinct genomic properties and separate clearly from all remaining chromosomes in the PCA built from chromosomal features (Supplementary Figure 4). The recovered microchromosomes showed elevated GC content and most were extremely repeat-rich relative to larger chromosomes (Figure 3A, Figure 3B). They contained few genes in absolute terms and remained gene-sparse even after accounting for their size (Figure 3C, Figure 3E). Despite their sparse gene content, most of their annotated genes were absent from the prior assembly (GCF_018691265.1, Figure 3D, Figure 3E).

### Haplotype synteny and a complex rearrangement on chromosome 7

Whole-genome alignment between haplotypes indicated extensive collinearity punctuated by a single, large and complex rearrangement on chromosome 7 (Figure 2A). Nearly the entire paternal chromosome 7 aligned in the opposite orientation to the maternal chromosome 7, with nested structure: a large (∼20 Mb) inversion interleaved with two shorter inverted segments and separated by two collinear blocks, all translocated relative to each other (Figure 4A). Repeat density increased near inferred breakpoints in both haplotypes, supporting the potential evolution of successive rearrangements (Figure 4a) [55].

**Figure 4.**
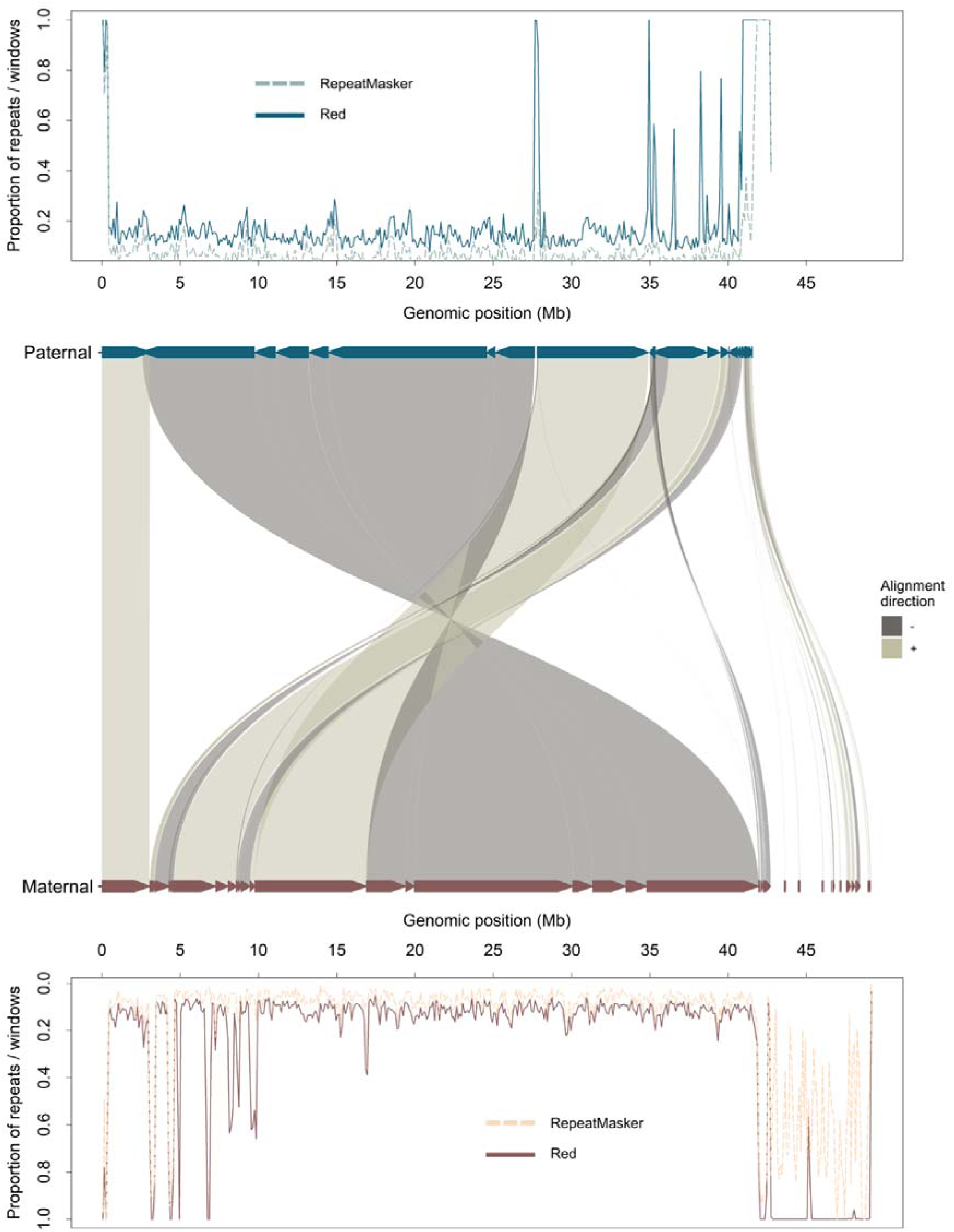
Large complex inversion on chromosome 7. Alignment between the paternal (top, dark blue) and maternal (bottom, burgundy) chromosome 7 highlights a major chromosomal inversion. The figure shows an extract from the whole-genome alignment, restricted to alignment blocks that are exclusive to chromosome 7 in both haplomes. Regions aligning in the same orientation are displayed in light grey, whereas regions aligned in the opposite orientation are shown in dark grey. Repeat proportion, calculated in 100-kbp windows, is plotted along the top and bottom. Two repeat detection methods are shown: RepeatMasker, represented in a lighter colour with a dashed line, and Red, represented in a dark colour with a solid line.

## Discussion

The telomere⍰to⍰telomere (T2T) era has transformed genome assembly, making it possible to reconstruct complete chromosomes across an increasing range of species [1–5,8,11]. Yet birds remain among the most challenging vertebrates to assemble owing to their numerous, small, GC-rich and repeat⍰dense microchromosomes, which have historically been under⍰represented or mis⍰assembled in reference genomes [7,23,24]. Here, we present the first haplotype⍰resolved, chromosome⍰level assemblies for an owl, the Western barn owl (*Tyto alba*), each comprising 46 chromosomes as expected from the karyotype [31,56] and substantially improving upon previous barn owl references [32,33]. These two haplotype-resolved assemblies exhibit high contiguity and completeness (with few gaps and BUSCO scores above 99%) and recover six chromosomes that were absent from earlier assemblies, providing the first near⍰complete representation of the species’ chromosomal assembly.

### A multi-layered assembly strategy

No single sequencing technology was sufficient to resolve the barn owl genome. Instead, the assembly succeeded because multiple complementary resources were integrated, with each compensating for the others’ blind spots. First, we phased the haplomes using Illumina reads from the parents, partitioning offspring long reads with trio aware *k*-mers before assembly to reduce switches and produce cleaner haplotype-specific contigs for downstream scaffolding. Then, HiFi contigs formed the structural backbone of the assembly, providing the accuracy and read length necessary to produce long, high⍰confidence unitigs. However, Hi-Fi reads alone cannot reliably span extremely GC⍰rich or repeat⍰dense regions, features abundant on avian micro- and dot chromosomes [57]. ONT ultra⍰long reads filled this gap, traversing repetitive sequences and bridging discontinuities between HiFi contigs, consistent with earlier reports demonstrating their unique value for difficult regions in bird genomes [25,26]. For chromosomes represented in the linkage map [34], population⍰based marker order provided a scaffold for organising HiFi / ONT contigs into linkage groups, after which Hi⍰C contact data confirmed chromosome⍰scale structure for the larger elements. However, Hi⍰C proved uninformative for microchromosomes: their short length, high repeat content, and poor mappability limit reliable detection of meaningful contact patterns, a difficulty also encountered in other avian assemblies [24]. To identify and assemble these highly challenging chromosomes, we instead relied on conserved microchromosome gene⍰set signature [24]. This allowed to pinpoint unplaced unitigs enriched for microchromosome⍰specific genes. These fragments were then manually curated and stitched together, guided by ONT long⍰read alignments that provided the only long⍰range continuity through repetitive landscapes.

Altogether, the assembly demonstrates that the integration of all the available information was critical to construct a robust, near⍰complete chromosomal assembly. Our recommendation to facilitate good future assemblies would be to collect enough material to generate several HiFi libraries of different fragment sizes and collect enough ONT sequences, preferably as long as possible, before starting the assembly. Filtering of sequences containing high number of internal repeats is also essential to prevent assembly artefacts.

### Recovery and characteristics of the missing chromosomes

Bird genomes remain difficult to assemble fully, and only a handful of avian references currently include the entire set of microchromosomes [23,25,26,28]. However, synteny⍰based scaffolding is often insufficient for microchromosomes because they show rapid evolution and poor conservation of gene order across species [23,24].

We assembled six previously missing chromosomes for the barn owl, displaying hallmark features of micro / dot chromosomes (elevated GC content and enriched repeats) consistent with other avian genomes [25,26,28]. However, in contrast with the micro-chromosomes present in the previous assembly, they remain gene⍰sparse relative to expectations based on the classic view of microchromosomes as strongly gene⍰dense [23,24]. This pattern may reflect genuine biological variation in the barn owl lineage but also residual annotation challenges in small repeat⍰rich elements now referred to as dot chromosomes [58].

Based on our feature□based analysis, the distinction between macro□, micro□, and dot□chromosomes is not straightforward, and no single variable (e.g., length, GC content, or gene content) provides clear boundaries between these chromosomal categories. Future work should examine the functional significance of genes located on these newly identified chromosomes and explore whether particular gene families (such as immunity□or sensory□related genes) exhibit micro□/ dot□chromosome–specific organization in owls.

### Phased assemblies reveal complex structural variation

Fully phased haplotypes allow the precise characterisation of complex structural variation. Structural variants (SVs), such as large inversions, are increasingly recognised as important drivers of adaptation [59], phenotypic evolution [60], and recombination landscape heterogeneity [61], but their detection and interpretation require high⍰quality, often phased reference genomes [7,8,55,62]. In barn owls, previous indirect evidence suggested the presence of a large inversion linked to climate [63], but its structure and extent could not be determined with pseudo⍰haploid or fragmented references [32,33]. Our phased assemblies reveal a ∼38 Mb complex inversion on chromosome 7, comprising multiple nested inverted and collinear segments. Yet the evolutionary history and potential phenotypic consequences of this inversion remain unknown. An important next step will be to examine its population frequency, heterozygosity patterns, and possible associations with traits such as coloration, dispersal, or behaviour traits that have shown geographic structure in barn owls [32].

### Conclusions and perspectives

Together, our phased, chromosome⍰level assemblies provide the most complete and accurate genomic resource to date for the barn owl and, more broadly, for the order Strigiformes [30,32,33]. They reveal previously unknown chromosomes, expose the architecture of a complex chromosomal inversion, and shed new light on microchromosome organisation in a lineage with a complex karyotype. These resources create a strong foundation for investigating structural variation, recombination dynamics, and the genetic basis of phenotypic divergence in barn owls. When combined with long-term ecological and population-genomic datasets available for this species, they will enable new levels of biological inference and contribute to a broader understanding of avian genome evolution.

## Material and methods

### Sampling

To build our genome assemblies, we used the same reference individual as in Machado et al. (2022) [32], a Swiss male barn owl (M040663), and its two parents (Father ring ID: M032330; Mother ring ID: M028977). On the 9^th^ of September 2019, we collected brachial-vein blood from each bird at a nestbox near Estavayer-le-Lac, Switzerland (46.85 N, 6.86 E). Blood samples were either kept at 4°C and process rapidly for High Molecular Weight (HMW) DNA extraction with agarose plugs (see below) or snap-frozen in liquid nitrogen and transferred to −80 °C freezer for long term preservation.

### High Molecular Weight DNA extraction of the reference individual

HMW DNA for the reference individual was extracted using an agarose plug method based on Zhang et al. [35] and previously described in Machado et al. (2021): fresh washed red blood cells were embedded in low-melt agarose plugs, digested with proteinase K, washed (EDTA; then 10 mM Tris pH 8.0, 1mM EDTA), melted, and digested with β-agarase to release the DNA. The resulting HMW DNA was dialysed, kept at 4 °C for 24 h, gently homogenised, quantified with Qubit (Thermo Scientific), and sized on Femto Pulse and pulsed-field gel electrophoresis. Mean fragment DNA size was around 100 to 150kbp. These DNA was used for PacBio HiFi SMRTbell Express 2.0 and for one ONT library.

To increase DNA yield, we extracted new HMW DNA from the −80°C frozen blood using MagAttract kit (Qiagen, elution repeated up to 4x to get enough DNA) for future PacBio library preparation. Frozen blood was also used to prepare HMW DNA with the Monarch® HMW DNA Extraction for Tissue kit (New England Biolabs) for future ONT libraries.

### Library preparation and sequencing

#### PacBio HiFi

The PacBio HiFi SMRTbell libraries were prepared and sequenced at the Lausanne Genome Technologies Facility (GTF, University of Lausanne). Briefly, HMW DNA was sheared (Megaruptor; Diagenode, Denville, NJ, USA) to ∼20 kb fragments and checked on a Fragment Analyser (Advanced Analytical Technologies, Ames, IA, USA). We prepared three SMRTbell libraries, two with SMRTBell Express 3.0 (10 µg DNA each) and one with Express 2.0 (5 µg), in accordance with the manufacturer’s recommendations. After BluePippin size selection (>15 kb; Sage Science, Inc., Beverly, MA, USA), libraries were sequenced on one and two SMRT 8M cells, respectively, using v2.2/v2.0 chemistry on a PacBio Sequel II instrument (Pacific Biosciences, Menlo Park, CA, USA) for a film run time of 30 hours and a pre-extension time of 120 minutes.

#### Oxford Nanopore Technology

Libraries were prepared from Monarch-HMW and plug-derived DNA using the ligation sequencing kit (SQK-LSK114). Sequencing was performed on PromethION flow cells (FLO-PRO114M) using a PromethION 2 Solo instrument (Oxford Nanopore Technologies, Oxford, UK) at the Gene Expression Core Facility (GECF, EPFL, Switzerland). For Monarch-HMW DNA, sequencing and base calling were carried out using MinKNOW software (v23.07.5) with the integrated Guppy base caller (v7.0.9) employing the super-accuracy (SUP) model. Plug-derived DNA libraries were sequenced using MinKNOW software (v25.03.9), and base calling was performed with Dorado (v1.0.0) using the SUP model, including duplex read analysis.

### Hi-C crosslinking

Without thawing, we diluted 10 µL of −80°C frozen blood from the focal individual in 926.9 µL of phosphate⍰buffered saline (PBS) and mixed gently. We then added 63.1 µL of 10% formaldehyde and incubated 20 minutes at room temperature (mixing periodically). We quenched with glycine (final concentration of 125 mM), followed by a 15⍰minute incubation at room temperature. We stored the sample at −80 °C and sent frozen to Phase Genomics (United States) for Hi⍰C library preparation. Sequencing was performed at the GTF (University of Lausanne, Switzerland).

### DNA extraction and sequencing of the parents

Following Cumer et al. (2025) [36], we extracted genomic DNA from whole blood with the DNeasy Blood & Tissue kit (Qiagen, Hilden, Germany). We prepared TruSeq DNA PCR-free libraries (Illumina) and sequenced multiplexed libraries on a HiSeq 2500 (PE high throughput) at the GTF (University of Lausanne, Switzerland).

### Identification of repeated sequences and HiFi read classification

To reduce assembly artefacts from extreme repeats, we first counted 64□mers in HiFi reads with Jellyfish v2.3.0 (-m 64 -C -c 3 -t 32 -L 50 -s 9,717,986). There were 1,793,627 64□mers occurring >1,000x. These 64-mers were offsets of the same patterns and carried several instances of smaller sub-motifs. For example, the six most frequent k□mers (each, >1.5 M) carried several instances of the sub□motif GTCACGTGGTC. The next four (each > 1.3 M) carried TTAGGGTTAGGGTTAGGG. We further inspected reads containing highly repeated motifs and noticed some were covered by multiple instances of identical sub-motifs, which we decided to filter prior to the assembly. Based on these observations, we placed HiFi reads into five categories based on the presence of the following motifs scanned in forward and reverse complement orientation:

- *Centromere⍰like*, characterised by at least 1 occurrence of the sequence CAGCACTGTTTCCTCCAAACTACCATGCTGCGCATAGCAATCCCTGTCTAGCTGCTTCCTTTCACTGGGAACACA, and labelled as *centromeric-like* due to the high conservation of a 162pb long repeat, compatible with a centromeric sequence [37]. This classification was further confirmed by the FISH experiment, see the *Karyotyping and FISH of telomeric and centromeric sequences* section below,
- *Kinetochore⍰like / pericentromeric repeat,* characterised by at least 3 occurrences of the sequence of the motif GTCACGTGGTC, labelled as such since these repeats are frequently found at extremities of reads composed of centromere-like repeats.
- *Telomere⍰like,* characterised by at least 3 occurrences of the motif TTAGGGTTAGGGTTAGGG, composed of multiple repeats of the vertebrate telomeric motif (TTAGGG) [38],
- *Microsatellite*, characterised by at least 1 occurrence of the sequence ACCTTCCCCCCACCCCTCCCTTCTTCCCGGGCTCAGCTCTGCTCCCGATATCTCTACCTCCTCC and labelled as *microsatellite* since this sequence has been previously described and was used to genotype microsatellite markers in the barn owl [39],
- Other (none of the above).

We excluded centromere⍰like and kinetochore⍰like reads since they contained repeated motifs spanning entire reads of length over 20kb but retained telomere⍰like (since they are at the end of chromosomes and less likely to confuse the assembler), and microsatellite⍰like whose repeated motif was present only in parts of the reads. This yielded a total of 7,826,051 retained HiFi reads, that were subsequently sorted by parental allele (see next section).

### Allele sorting of HiFi reads

With the final intention to build haplome specific assembly, we sorted HiFi reads according to the parent of origin using a k-mer based approach that identifies parent-specific kmers in each read.

We first build parent-specific sets of k-mers from parental Illumina reads. To balance uneven parental sequencing depths (Father 172’731’951 reads vs Mother 93’360’939 reads) and sampling bias of kmers during sequencing, we randomly drew non-overlapping subsets of 46,680,469 Illumina paired-end reads, 3 for the Father and 2 for the Mother. In each of these five sets, we counted 49-mers (length chosen to avoid palindromes and reduce homopolymer errors; Supplementary Table 4), free of Ns. Finally, we defined nine parent specific k-mer libraries containing 49-mers present ≥ 2x within a parental subset and absent from pairs of subsets of the other parent (see Supplementary Table 5). These parent specific k-mer libraries allowed to account for sampling bias and statistical testing during the read classification procedure described in the next paragraph. This comparison between parental sets was done using a modified jellyfish v2.3.0 to emit k-mer “bin” files unique to one library and absent from a given set of other libraries (patch provided at: https://github.com/hugocrvl/TytoAlba_NewReferenceGenome/blob/main/2_AlleleSorting/Jellyfish-2.3.0-patch.txt).

We sorted the HiFi reads of the reference individual into two parental sets. Each HiFi read was scanned with a sliding window of 49□mers, using a 7□nt step to keep the computational burden manageable. For each read, we counted the occurrences of parent□specific k□mers. This counting procedure was performed for every pairwise combination of the parent□specific k□mer libraries, generating a distribution of k□mer counts for each parent for each read. Based on these distributions of counts, we assigned reads to parental origin using paired t□tests with Benjamini–Hochberg FDR control implemented in R. Reads with FDR ≤ 0.001 were assigned to the parent with the highest number of matching k□mers. Unassigned reads were rescanned using a 1□nt step sliding window, and the assignment test was repeated using an FDR threshold of ≤ 0.05. FDR control was performed separately within each read class (telomere□like, microsatellite□like, other) to account for compositional biases associated with these sequence categories (Supplementary Table 6).

### ONT reads filtering, classification and allele sorting

For ONT reads, we selected all reads generated by the ONT caller that were 25k nucleotides or longer. ONT reads were then attributed to the five categories described earlier (*Centromere⍰like, Kinetochore⍰like / pericentromeric repeat, Telomere⍰like, Microsatellite, other*) using the motifs identified with HiFi reads. The allelic sorting process explained in the previous section was applied to the ONT reads 1 and 2 (≥ 25-kbp long) using a sliding window offset of 7nt and an adjusted p-value ≤ 0.05 to assign predicted parental origin. This yielded a total of 1,024,293 retained ONT reads. Detailed results after the filtering, classification and sorting are presented in Supplementary Table 7.

### Haplotype-specific assembly and scaffolding

#### HiFi-based assemblies

We ran hifiasm 0.19.8-r603 to build separate maternal and paternal haplotype using: (i) HiFi reads assigned to that parent, (ii) all HiFi reads from uncertain origins and (iii) ONT from run 1 (ONT run 2 not available at that stage). After the initial assemblies and to detect regions with abnormally high parental coverage, we realigned HiFi reads to the draft assemblies with minimap2 2.26-r1175. After realigning the reads to both parental haplotypes, we assigned each read to the parent with the highest alignment score. After that step we identified regions where the coverage was <=20% and >=80% to the expected coverage. Reads contributing to these imbalances were identified, sorted by increasing adjusted p-value from the read sorting step (see above for details) and the bottom half of them re-assigned to the other parental allele, implementing the idea that the increased coverage is due to wrongly assigned reads. After this final reclassification of 14,717 reads, we generated two final haplotype assemblies using hifiasm 0.19.8-r603. The final drafts comprised 287 paternal and 331 maternal unitigs.

#### ONT assemblies

Using hifiasm 0.25.0-r726, we independently assembled ONT reads into parental haplotype (inputs: ONT runs 1 and 2 assigned to that parent + uncertain-origin ONT reads from runs 1 and 2), yielding 1030 paternal and 985 maternal unitigs

#### Final scaffolding with linkage groups and ONT-based assemblies

We first scaffolded the HiFi-based contigs into 40 linkage groups using the linkage map produced by Topaloudis et al. [34]: we extracted ±100 bp around each marker from the old assembly to create pseudo-reads and mapped them to the new haplotype assemblies, thus ordering unitigs by marker order and coordinates.

Then, we mapped all HiFi reads to their HiFi unitigs to flag low coverage regions of interest (ROI). Using ad-hoc scripts and aligners (minimap2, LASTZ, MAFFT), we stitched HiFi and ONT unitigs across ROIs and unitig ends guided by the linkage map, to produce the final scaffolds for both haplotypes. Because owl chromosomes are acrocentric (Figure 2B) [31,40], and we excluded centromere-rich reads, we oriented all scaffolds with a 5’ telomere (e.g.,5p-CCCTAA-3p repeat) at the p arm so that adding centromeres later will not alter the coordinate system.

### Recovery of micro-chromosomes

#### Identification of the microchromosomes

To recover the unscaffolded microchromosomes in both assemblies, we used MicroFinder [24]. This tool compares the gene content of unplaced scaffolds against a curated set of genes known to be enriched on microchromosomes across avian species, flagging candidates showing significant enrichment. Following the authors’ recommendations, we set a maximum scaffold size cutoff of 5 Mb to focus the analysis on short, genuinely unplaced sequences and retrieved 14 and 18 micro-chromosome contigs for the paternal and the maternal assemblies, respectively.

#### Assembly of the microchromosomes

Candidate unitigs identified in the *Recovery of microchromosomes* section were then manually assembled into six additional microchromosomes. This manual curation was done using ad-hoc scripts and aligners (minimap2, LASTZ, MAFFT) and supported by sequence alignment comparisons between the maternal and paternal haplotypes as well as using the unitigs produced by HiFi and ONT assemblies.

### Detection of terminal features of the chromosomes

We assessed whether chromosomes were assembled completely (i.e., spanning telomere□to□centromere for autosomes or telomere□to□telomere for the Z chromosome). Because centromeric and pericentromeric reads were excluded during assembly, we realigned the raw long-read data to both the maternal and paternal assemblies to identify reads mapping to chromosome ends and containing telomeric or centromeric/pericentromeric repeats. Both PacBio HiFi (Sequel IIe, chemistry M64) and Oxford Nanopore Technologies long reads were mapped against each reference. Each read was then assigned to a genomic feature class (telomere, centromere, kinetochore, microsatellite repeat, or “other”) based on the presence of class□specific sequence motifs (see Identification of repeated sequences and HiFi reads classification). Only primary alignments were retained (filtering on tp:A:P), and a per□read mapping proportion was computed as the number of matched bases divided by the total read length.

Reads mapping near chromosome termini (where telomeric and centromeric repeats are expected) were identified by selecting those with alignment start positions within 10 kb of the chromosome beginning or alignment end positions within 10 kb of the chromosome end. Terminal□proximal reads were further filtered to retain only those with mapping quality >10 and mapping proportion >5%. Finally, reads not classified as telomeric, centromeric, or kinetochore-like were removed.

Finally, reads were grouped by parental haplotype, chromosome, terminus (start or end), and feature class. Clusters supported by fewer than 15 reads were discarded. Because all scaffolds were oriented based on the presence of telomere□specific repeats (e.g., 5□CCCTAA□3′), we then examined the distribution of read clusters along each chromosome. Specifically, we inspected autosomes for clusters of telomeric reads at their chromosome starts, and the Z chromosome for telomeric clusters at both termini. Conversely, we assessed chromosome ends for the presence of centromeric or pericentromeric read clusters, consistent with expectations for scaffold boundaries corresponding to centromeric regions.

### Protein-based synteny across avian species

To confirm the identification of the Z chromosome, we aligned chicken (GCF_016699485.2) and zebra finch (GCF_048771995.1) proteins to both haplotype assemblies using miniprot (v0.18-r281) [41]. We kept alignments with ≥ 70% sequence identity and ≥ 90% target coverage, then merged pairwise mapping formats (PAFs) with reference feature tables to count matches per reference chromosome vs barn owl chromosome. Sankey diagrams summarised chromosome-level correspondences (Supplementary Figure 1 and 2).

### Evaluation of the phasing of the assemblies

To avoid biases caused by differences in parental sequencing depth, we down-sampled parental Illumina reads to 80,000,000 per parent using seqtk (v1.4) with a fixed seed (-s 100) for reproducibility and counted 49⍰mers with meryl *count* (v1.4.1). We derived parental⍰private k⍰mers (meryl *difference*), counted 49-mers in each assembly, and compute a phasing error as: the number of misplaced parental-private k-mer in the wrong haplotype divided by the total number of parental-private k-mers across both haplotypes.

### Genome quality control

Quality and completeness were evaluated at two stages of the assembly process: the contig stage (prior to scaffolding) and the final parental assemblies. We assessed completeness with BUSCO v6.0.0 (aves_odb10) [42]. We compute the general assembly statistics (contig/scaffold counts, N50/N90, GC, gaps) with Quast v5.3.0 [43] and visualised with BlobTools2 snailplots [44].

### Repeat annotation

We identified the repeats with two complementary approaches: (i) Red (de novo, library free) [45] and (ii) RepeatModeler v2.0.4 [46] with default parameters to build a custom repeat library, followed by RepeatMasker v4.1.5 [47] with default parameters for classification and masking of the repeats.

### Gene annotation

We annotated both haplomes with NCBI Eukaryotic Genome Annotation Pipeline (EGAPx) [48]. In brief, EGAPx takes a genome assembly in FASTA format, a taxonomy ID, and RNA-seq data as input. First, it uses protein homology by aligning protein sequences from closely related species with prosplign. Second, it uses transcriptome information from the focal species. We included RNA-seq short reads from six tissues (liver, heart, kidney, testis, growing back feathers, and thalamus) previously published in Ducrest et al. [33], which EGAPx aligns with STAR [49] and minimap2 [50]. Third, EGAPx runs Gnomon to generate *ab-initio* predictions [51]. These different hits are then integrated by Gnomon to produce a final consensus annotation.

We realigned Coding DNA Sequence (CDS) from the EGAPx annotation to their respective haplome with minimap2 in split mode [50] and kept only full-length mapping, MAPQ = 60. These quality-filtered CDS were then mapped back to the previous reference genome (T.alba_DEE_v4.0 - GCF_018691265.1) to classify CDS as previously present (full-length, MAPQ = 60) or new. For multi-mapped CDS, we retained the alignment with the longest aligned span and summarised counts per chromosome.

### Chromosome characteristics and classification

To distinguish macro-, micro-, and dot chromosomes, we ran a PCA on z-scored, chromosome-level features: (i) length, (ii) total exonic length (relative), (iii) GC, (iv) gene density, (v) exon density, (vi) mean exon length; (vii) fraction of simple/low complexity repeats and (viii) fraction of other RepeatMasker classes (resolving overlaps by the larger element’s class).

### Genome mappability masking

Genome-wide mappability was quantified using the seqbility/SNPable pipeline (available at https://github.com/lh3/misc/tree/master/seq/seqbility). The reference genome was first decomposed into all possible k-mers using the splitfa program, with the k-mer size of 150bp, matching common read length. These k-mers were then aligned back to the reference using BWA in aln mode [52]. To match the SNPable specification, we used several non□default BWA parameters to allow permissive and sensitive mapping: the seed length was reduced (-l 32), up to two seed mismatches were permitted (-k 2), and both the gap□open (-O 3) and gap□extension (-E 3) penalties were increased relative to standard BWA defaults.

The resulting SAM file was processed with gen_raw_mask.pl, which extracts, for each k-mer, the number of perfect and near⍰perfect alignments it produces across the genome. This raw mask was then summarised into position⍰wise mappability tracks using the gen_mask script. For this step several non⍰default stringency thresholds were applied (-r 0.50, 0.90, 0.95, and 0.99). These thresholds represent the minimum fraction of overlapping k-mers that must map uniquely for a genomic position to be considered mappable. Lower thresholds generate more permissive callable regions, whereas higher thresholds identify only the most confidently unique regions.

### Recombination map

We lifted the coordinates of the linkage markers from [34] by extracting 200-bp around each SNP with bedtools *getfasta* (v2.x) and mapping these sequences to each haplome (bwa⍰mem v0.7.x, sorted with samtools v1.x). We kept only SNPs with a unique and perfect alignments (MAPQ = 60), and for each combined the new physical coordinates with the original linkage⍰map metadata (linkage group and genetic position) to lift the recombination maps coordinates in each haplotype coordinate system.

### Hi-C contact maps

We sequenced 338,385,433 Hi-C pairs. After enzyme-site filtering for GATC at R1 ends and R2 starts, 112,239,344 paired reads (33.2%) remained. We aligned reads with FetchGWI [53], converted alignments to genomic pairs, and built .cool files with cooler v0.9.3 (cooler *cload pairs* c1 1 -p1 2 -c2 3 -p2 4), followed by cooler *zoomify*. Genome-wide maps were generated at 16-kb -wide bins.

### Karyotyping and FISH of telomeric and centromeric sequences

#### Cell culture and chromosome preparation

On the 10^th^ of July 2025, we collected three growing breast feathers from a 28-day-old female nestling (M046151). After surface sterilisation (70% ethanol) and PBS rinses, we dissected the feather basal parts, scraped inner tissues using tweezers, added it to a 24-well plate containing Dulbecco’s Modified Eagle’s Medium high glucose (DMEM, Avantor) with 20% foetal bovine serum (FBS, Sigma), 1 % penicillin-streptomycin (Thermo Scientific), 5% AmnioMax II (Gibco, Thermo Scientific) and the MycoGenie MycoPlasma Elimination Kit (1:2000; Assay Genie, Chemie Brunschwig). Cultures were maintained at 37°C with 5% CO_2_ for ≥ 3 weeks to obtain enough cells.

To arrest mitosis, we treated cells with KaryoMax Colcemid (0.5 µg/ml, 1h; Thermo Scientific), followed by 0.075 M KCl hypotonic shock (20 minutes, 37°C) and gentle fixation in fresh methanol/acetic acid (3:1) with 3-4 wash/fix cycles. The metaphase chromosome suspensions were stored at −20°C.

#### Hybridisation

We spread 10 µl of metaphase chromosome suspension per slide. After aging (1h, 65°C) and a 2x SSC wash (10 minutes, 300 mM sodium chloride, 30 mM sodium citrate, pH 7.0) at room temperature, slides were incubated with 100 µg/ml RNAse A (Qiagen) in 2x SSC for 45 minutes at 37°C. The slides were then washed in 2x SSC for 5 minutes at room temperature and treated with 0.005% pepsin for 3 minutes in 0.01 N HCl heated to 37°C. We then washed in PBS with 50 mM MgCl₂, denatured them in 1% paraformaldehyde (Thermo Scientific) for 10 minutes at room temperature, washed with 2x SSC, and dehydrated using a series of 70%, 90%, and 100% ethanol and let them dry for 20 minutes at room temperature. We diluted a centromeric probe (1 μl of 10 µM, CEN3.2_ATTO590, 3’-TTTCCTCCAAACTACCATGCTGCG-5’, see the *Centromere*⍰*like* section above for the identification of this sequence) in 20 μl containing the PNA telomere hybridisation buffer with 10 μl of Telomere PNA FITC probe (Telomere PNA Kit/FITC for Flow Cytometry, PNA FISH Kit/FITC, Agilent Technologies). The mixture was applied to the chromosome spread. Coverslips were sealed, slides denatured (92°C, 3 minutes) and hybridised overnight (37°C). The slides were then washed twice in 2x SSC + 0.1% Tween 20 (60°C, 15 minutes), and once in 0.2x SSC (room temperature), then mounted with Vectashield mounting medium + DAPI (500 ng/ml; Sigma). Imaging used a Leica Stellaris5 confocal microscope equipped with a 63xPlanApo oil immersion objective (CIF platform, University of Lausanne). All acquisitions were produced by superimposing focal planes. Post-processing, including pseudo colouring, was performed using Fiji [54].

## Authors contribution

H.C.: conceptualisation, data curation, formal analysis, validation, investigation, visualization, methodology, writing (original draft; review & editing). M.B.S.: data curation, formal analysis, visualization, writing (review & editing). A.B.: data curation, formal analysis, visualization, writing (review & editing). A.T.: data curation, writing (review & editing). C.S.: Resources, writing (final revision). E.C.: Resources, writing (final revision). D.C.: Resources, writing (final revision). B.A.: Resources, funding acquisition, writing (final revision). A.R.: Resources, funding acquisition, writing (final revision). A.L.D.: conceptualisation, resources, formal analysis, investigation, visualization, methodology, writing (review & editing). N.G.: conceptualisation, data curation, software, formal analysis, validation, investigation, visualization, methodology, writing (review & editing). C.I.: conceptualisation, data curation, software, formal analysis, validation, visualization, methodology, writing (review & editing). T.C.: conceptualisation, data curation, formal analysis, supervision, validation, investigation, visualization, methodology, writing (original draft; review & editing). J.G.: conceptualisation, supervision, funding acquisition, methodology, writing (review & editing)

## Data availability

Samples were collected under Swiss veterinary permit 35712, VD3884. The data underlying this article are available under the BioProjects PRJNA440222 and PRJNA440227. FASTA sequences, repeat annotations, gene annotations, recombination maps, and mappability masks for both haplotypes are available at Zenodo: https://zenodo.org/records/19111812?token=eyJhbGciOiJIUzUxMiIsImlhdCI6MTc3MzkzMTI 2NCwiZXhwIjoxNzk5NTM5MTk5fQ.eyJpZCI6IjJkMWY3N2Q0LTc2YjEtNDIzZi1hODU4LTA1 ZTBhNTM5Zjk0YyIsImRhdGEiOnt9LCJyYW5kb20iOiIzMGMxZmYzZDNhOGU4OGY2MDNl MmJmNTNhMzAxZjdhYSJ9.8ghGfu1RQQd1I8w3BvX4L_FI8CUcyQ3aYHAQiLg23anne_D3i FHz5R2pOdtGY52SNIgPCW29mS2471UwJxWdbQ. (note: this temporary link will be replaced with a permanent DOI upon publication).

The code used in this study is accessible at: https://github.com/hugocrvl/TytoAlba_NewReferenceGenome. Additional data or information can be obtained from the corresponding authors upon reasonable request.

## Acknowledgement

We thank the entire field team of the groups of Alexandre Roulin and Bettina Almasi for their contribution in collecting samples. In particular, we are grateful to Roman Bühler, Nicolas Sironi, Kim Schalcher, Nathan Külling, Alexandre Roland, and Ilou Lejeune for collecting the blood and feather samples used in this study. We also acknowledge the Genome Technologies Facility (GTF) at the University of Lausanne and the Gene Expression Core Facility (GECF) at EPFL for their technical guidance and expertise during sequencing.

## References

1. Nurk S, Koren S, Rhie A, Rautiainen M, Bzikadze AV, Mikheenko A, et al.. The complete sequence of a human genome. Science. 2022; doi: 10.1126/science.abj6987.

2. Chen J, Wang Z, Tan K, Huang W, Shi J, Li T, et al.. A complete telomere-to-telomere assembly of the maize genome. Nat Genet. 2023; doi: 10.1038/s41588-023-01419-6.

3. Shang L, He W, Wang T, Yang Y, Xu Q, Zhao X, et al.. A complete assembly of the rice Nipponbare reference genome. Molecular Plant. 2023; doi: 10.1016/j.molp.2023.08.003.

4. Wu H, Luo L-Y, Zhang Y-H, Zhang C-Y, Huang J-H, Mo D-X, et al.. Telomere-to-telomere genome assembly of a male goat reveals variants associated with cashmere traits. Nat Commun. 2024; doi: 10.1038/s41467-024-54188-z.

5. Yoo D, Rhie A, Hebbar P, Antonacci F, Logsdon GA, Solar SJ, et al.. Complete sequencing of ape genomes. Nature. 2025; doi: 10.1038/s41586-025-08816-3.

6. Belton J-M, McCord RP, Gibcus JH, Naumova N, Zhan Y, Dekker J. Hi-C: a comprehensive technique to capture the conformation of genomes. Methods. 2012; doi: 10.1016/j.ymeth.2012.05.001.

7. Zhang T, Zhou J, Gao W, Jia Y, Wei Y, Wang G. Complex genome assembly based on long-read sequencing. Briefings in Bioinformatics. 2022; doi: 10.1093/bib/bbac305.

8. Li H, Durbin R. Genome assembly in the telomere-to-telomere era. Nat Rev Genet. 2024; doi: 10.1038/s41576-024-00718-w.

9. Logsdon GA, Ebert P, Audano PA, Loftus M, Porubsky D, Ebler J, et al.. Complex genetic variation in nearly complete human genomes. Nature. 2025; doi: 10.1038/s41586-025-09140-6.

10. Mosbruger TL, Dinou A, Duke JL, Ferriola D, Li Y, Hayeck TJ, et al.. Haplotypic resolution of the challenging genomic regions of MHC and KIR using a combination of targeted sequencing and a novel assembly pipeline. Nucleic Acids Research. 2025; doi: 10.1093/nar/gkaf441.

11. Yang Y, Du W, Li Y, Lei J, Pan W. Recent advances and challenges in *de novo* genome assembly. GCOMM. 2025; doi: 10.48130/gcomm-0025-0015.

12. Koren S, Rhie A, Walenz BP, Dilthey AT, Bickhart DM, Kingan SB, et al.. De novo assembly of haplotype-resolved genomes with trio binning. Nat Biotechnol. 2018; doi: 10.1038/nbt.4277.

13. Cheng H, Concepcion GT, Feng X, Zhang H, Li H. Haplotype-resolved de novo assembly using phased assembly graphs with hifiasm. Nat Methods. 2021; doi: 10.1038/s41592-020-01056-5.

14. Selvaraj S, R Dixon J, Bansal V, Ren B. Whole-genome haplotype reconstruction using proximity-ligation and shotgun sequencing. Nat Biotechnol. 2013; doi: 10.1038/nbt.2728.

15. Cheng H, Jarvis ED, Fedrigo O, Koepfli K-P, Urban L, Gemmell NJ, et al.. Haplotype-resolved assembly of diploid genomes without parental data. Nat Biotechnol. 2022; doi: 10.1038/s41587-022-01261-x.

16. Falconer E, Lansdorp PM. Strand-seq: A unifying tool for studies of chromosome segregation. Seminars in Cell & Developmental Biology. 2013; doi: 10.1016/j.semcdb.2013.04.005.

17. Porubsky D, Ebert P, Audano PA, Vollger MR, Harvey WT, Marijon P, et al.. Fully phased human genome assembly without parental data using single-cell strand sequencing and long reads. Nat Biotechnol. 2021; doi: 10.1038/s41587-020-0719-5.

18. Chin C-S, Peluso P, Sedlazeck FJ, Nattestad M, Concepcion GT, Clum A, et al.. Phased diploid genome assembly with single-molecule real-time sequencing. Nat Methods. 2016; doi: 10.1038/nmeth.4035.

19. Low WY, Tearle R, Liu R, Koren S, Rhie A, Bickhart DM, et al.. Haplotype-resolved genomes provide insights into structural variation and gene content in Angus and Brahman cattle. Nat Commun. 2020; doi: 10.1038/s41467-020-15848-y.

20. Ebert P, Audano PA, Zhu Q, Rodriguez-Martin B, Porubsky D, Bonder MJ, et al.. Haplotype-resolved diverse human genomes and integrated analysis of structural variation. Science. 2021; doi: 10.1126/science.abf7117.

21. Rhie A, McCarthy SA, Fedrigo O, Damas J, Formenti G, Koren S, et al.. Towards complete and error-free genome assemblies of all vertebrate species. Nature. 2021; doi: 10.1038/s41586-021-03451-0.

22. Kong W, Wang Y, Zhang S, Yu J, Zhang X. Recent Advances in Assembly of Complex Plant Genomes. Genomics, Proteomics & Bioinformatics. 2023; doi: 10.1016/j.gpb.2023.04.004.

23. Peona V, Weissensteiner MH, Suh A. How complete are “complete” genome assemblies?—An avian perspective. Molecular Ecology Resources. 2018; doi: 10.1111/1755-0998.12933.

24. Mathers TC, Paulini M, Sotero-Caio CG, Wood JMD. MicroFinder: Conserved gene-set mapping and assembly ordering for manual curation of bird microchromosomes. Genomics;

25. Zhao Q, Yin Z, Hou Z. Near telomere-to-telomere genome assemblies of Silkie Gallus gallus and Mallard Anas platyrhynchos restored the structure of chromosomes and “missing” genes in birds. J Animal Sci Biotechnol. 2025; doi: 10.1186/s40104-024-01141-1.

26. Huang Z, Xu Z, Bai H, Huang Y, Kang N, Ding X, et al.. Evolutionary analysis of a complete chicken genome. Proc Natl Acad Sci USA. 2023; doi: 10.1073/pnas.2216641120.

27. Luo H, Jiang X, Li B, Wu J, Shen J, Xu Z, et al.. A high-quality genome assembly highlights the evolutionary history of the great bustard (Otis tarda, Otidiformes). Commun Biol. 2023; doi: 10.1038/s42003-023-05137-x.

28. Hu J, Song L, Ning M, Niu X, Han M, Gao C, et al.. A new chromosome-scale duck genome shows a major histocompatibility complex with several expanded multigene families. BMC Biol. 2024; doi: 10.1186/s12915-024-01817-0.

29. Degrandi TM, Barcellos SA, Costa AL, Garnero ADV, Hass I, Gunski RJ. Introducing the Bird Chromosome Database: An Overview of Cytogenetic Studies in Birds. Cytogenet Genome Res. 2020; doi: 10.1159/000507768.

30. Baalsrud HT, Garmann-Aarhus B, Enevoldsen ELG, Krabberød AK, Fischer D, Tooming-Klunderud A, et al.. Evolutionary new centromeres in the snowy owl genome putatively seeded from a transposable element.

31. Belterman RHR, De Boer LEM. A karyological study of 55 species of birds, including karyotypes of 39 species new to cytology. Genetica. 1984; doi: 10.1007/BF00056765.

32. Machado AP, Cumer T, Iseli C, Beaudoing E, Ducrest A, Dupasquier M, et al.. Unexpected post-glacial colonisation route explains the white colour of barn owls ( *Tyto alba*) from the British Isles. Mol Ecol. 2021; doi: 10.1111/mec.16250.

33. Ducrest A, Neuenschwander S, Schmid-Siegert E, Pagni M, Train C, Dylus D, et al.. New genome assembly of the barn owl ( *Tyto alba alba* ). Ecology and Evolution. 2020; doi: 10.1002/ece3.5991.

34. Topaloudis A, Cumer T, Lavanchy E, Ducrest A-L, Simon C, Machado AP, et al.. The recombination landscape of the barn owl, from families to populations. Wright S, editor. GENETICS. 2025; doi: 10.1093/genetics/iyae190.

35. Zhang M, Zhang Y, Scheuring CF, Wu C-C, Dong JJ, Zhang H-B. Preparation of megabase-sized DNA from a variety of organisms using the nuclei method for advanced genomics research. Nat Protoc. 2012; doi: 10.1038/nprot.2011.455.

36. Cumer T, Topaloudis A, Lavanchy E, Ducrest A-L, Cabello SS, Hewett A, et al.. Genomic bases of short-term evolution in the wild revealed by long-term monitoring and population-scale sequencing. Evolutionary Biology;

37. Melters DP, Bradnam KR, Young HA, Telis N, May MR, Ruby JG, et al.. Comparative analysis of tandem repeats from hundreds of species reveals unique insights into centromere evolution. Genome Biol. 2013; doi: 10.1186/gb-2013-14-1-r10.

38. Meyne J, Ratliff RL, Moyzis RK. Conservation of the human telomere sequence (TTAGGG)n among vertebrates. Proc Natl Acad Sci USA. 1989; doi: 10.1073/pnas.86.18.7049.

39. Burri R, Antoniazza S, Siverio F, Klein Á, Roulin A, Fumagalli L. Isolation and characterization of 21 microsatellite markers in the barn owl ( *Tyto alba* ). Molecular Ecology Resources. 2008; doi: 10.1111/j.1755-0998.2008.02121.x.

40. Rebholz WER, Boer LEMD, Sasaki M, Belterman RHR, Nishida-Umehara C. The Chromosomal Phylogeny of Owls (Strigiformes) and New Karyotypes of Seven Species. cytologia. 1993; doi: 10.1508/cytologia.58.403.

41. Li H. Protein-to-genome alignment with miniprot. arXiv;

42. Simão FA, Waterhouse RM, Ioannidis P, Kriventseva EV, Zdobnov EM. BUSCO: assessing genome assembly and annotation completeness with single-copy orthologs. Bioinformatics. 2015; doi: 10.1093/bioinformatics/btv351.

43. Gurevich A, Saveliev V, Vyahhi N, Tesler G. QUAST: quality assessment tool for genome assemblies. Bioinformatics. 2013; doi: 10.1093/bioinformatics/btt086.

44. Challis R, Richards E, Rajan J, Cochrane G, Blaxter M. BlobToolKit – Interactive Quality Assessment of Genome Assemblies. G3 Genes|Genomes|Genetics. 2020; doi: 10.1534/g3.119.400908.

45. Girgis HZ. Red: an intelligent, rapid, accurate tool for detecting repeats de-novo on the genomic scale. BMC Bioinformatics. 2015; doi: 10.1186/s12859-015-0654-5.

46. Flynn JM, Hubley R, Goubert C, Rosen J, Clark AG, Feschotte C, et al.. RepeatModeler2 for automated genomic discovery of transposable element families. Proc Natl Acad Sci USA. 2020; doi: 10.1073/pnas.1921046117.

47. Smit AFA, Hubley R. RepeatModeler Open-1.0.

48. NCBI. EGAPx: Eukaryotic Genome Annotation Pipeline – External. GitHub;

49. Dobin A, Davis CA, Schlesinger F, Drenkow J, Zaleski C, Jha S, et al.. STAR: ultrafast universal RNA-seq aligner. Bioinformatics. 2013; doi: 10.1093/bioinformatics/bts635.

50. Li H. Minimap2: pairwise alignment for nucleotide sequences. Birol I, editor. Bioinformatics. 2018; doi: 10.1093/bioinformatics/bty191.

51. Souvorov A, Kapustin Y, Kiryutin B, Chetvernin V, Tatusova T, Lipman D. Gnomon – NCBI eukaryotic gene prediction tool. 2020;

52. Li H. Aligning sequence reads, clone sequences and assembly contigs with BWA-MEM. arXiv;

53. Iseli C, Ambrosini G, Bucher P, Jongeneel CV. Indexing Strategies for Rapid Searches of Short Words in Genome Sequences. Wu X, editor. PLoS ONE. 2007; doi: 10.1371/journal.pone.0000579.

54. Schindelin J, Arganda-Carreras I, Frise E, Kaynig V, Longair M, Pietzsch T, et al.. Fiji: an open-source platform for biological-image analysis. Nat Methods. 2012; doi: 10.1038/nmeth.2019.

55. Gozashti L, Harringmeyer OS, Hoekstra HE. How repeats rearrange chromosomes: The molecular basis of chromosomal inversions in deer mice. Cell Reports. 2025; doi: 10.1016/j.celrep.2025.115644.

56. Renzoni A, Vegni-Talluri M. The karyograms of some Falconiformes and Strigiformes. Chromosoma. 1966; doi: 10.1007/BF00335203.

57. Lang D, Zhang S, Ren P, Liang F, Sun Z, Meng G, et al.. Comparison of the two up-to-date sequencing technologies for genome assembly: HiFi reads of Pacific Biosciences Sequel II system and ultralong reads of Oxford Nanopore. GigaScience. 2020; doi: 10.1093/gigascience/giaa123.

58. Hron T, Miklík D, Pačes J, Pajer P, Pečenka V, Hejnar J, et al.. Decoding the avian missing gene mystery: dot chromosomes unmask extensive gene loss and novel genetic instability. Genomics;

59. Lundberg M, Liedvogel M, Larson K, Sigeman H, Grahn M, Wright A, et al.. Genetic differences between willow warbler migratory phenotypes are few and cluster in large haplotype blocks. Evolution Letters. 2017; doi: 10.1002/evl3.15.

60. Küpper C, Stocks M, Risse JE, Dos Remedios N, Farrell LL, McRae SB, et al.. A supergene determines highly divergent male reproductive morphs in the ruff. Nat Genet. 2016; doi: 10.1038/ng.3443.

61. Faria R, Johannesson K, Butlin RK, Westram AM. Evolving Inversions. Trends in Ecology & Evolution. 2019; doi: 10.1016/j.tree.2018.12.005.

62. Mahmoud M, Gobet N, Cruz-Dávalos DI, Mounier N, Dessimoz C, Sedlazeck FJ. Structural variant calling: the long and the short of it. Genome Biol. 2019; doi: 10.1186/s13059-019-1828-7.

63. Corval H, Cumer T, Topaloudis A, Collart F, Guisan A, Roulin A, et al.. Genetically diverse populations hold the keys to climatic adaptation: a lesson from a cosmopolitan raptor.

