## Supplementary material for "A chromosome-level, haplotype-resolved genome assembly for the barn owl, *Tyto alba*": SupplementaryMaterial.docx

**Supplementary Figures**

**Supplementary Tables**

### Supplementary Figures

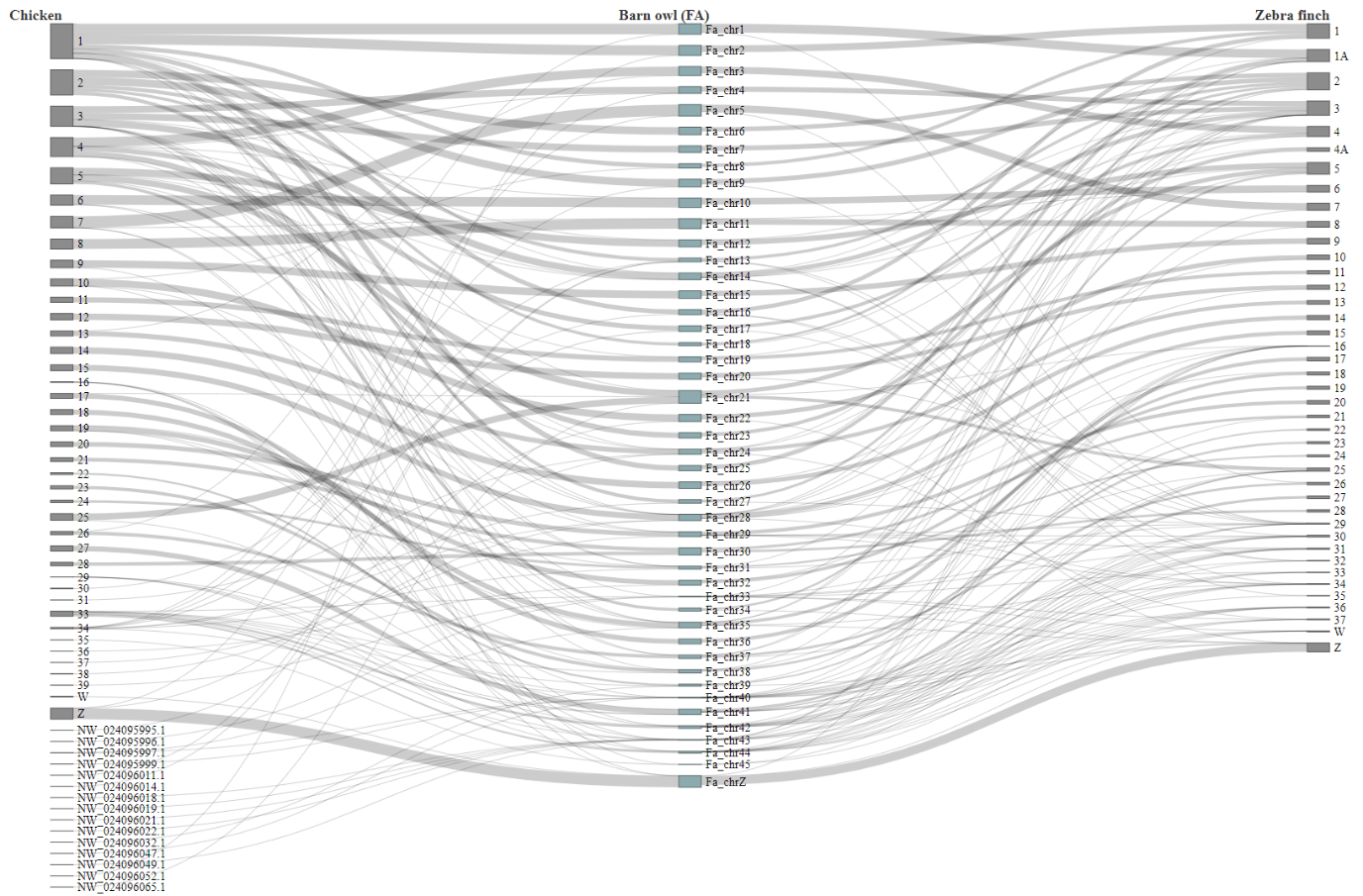

Figure S1

**Miniprot alignments for the paternal proteome.** Protein alignments of the chicken and zebra finch proteome against the paternal haplome (only matches with ≥ 70% sequence identity and ≥ 90% target coverage retained).

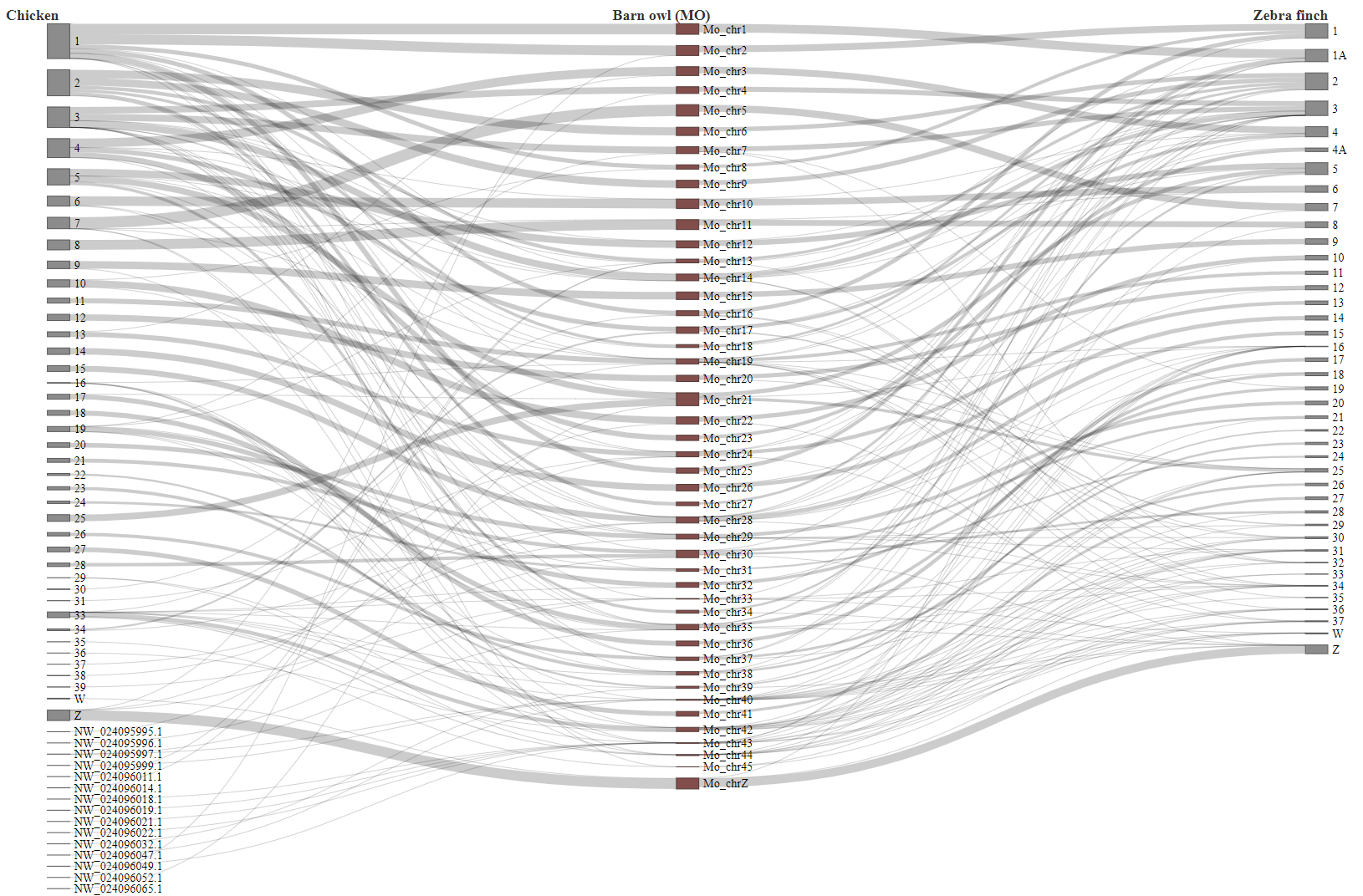

Figure S2

**Miniprot alignments for the maternal proteome.** Protein alignments of the chicken and zebra finch proteome against the maternal haplome (only matches with ≥ 70% sequence identity and ≥ 90% target coverage retained).

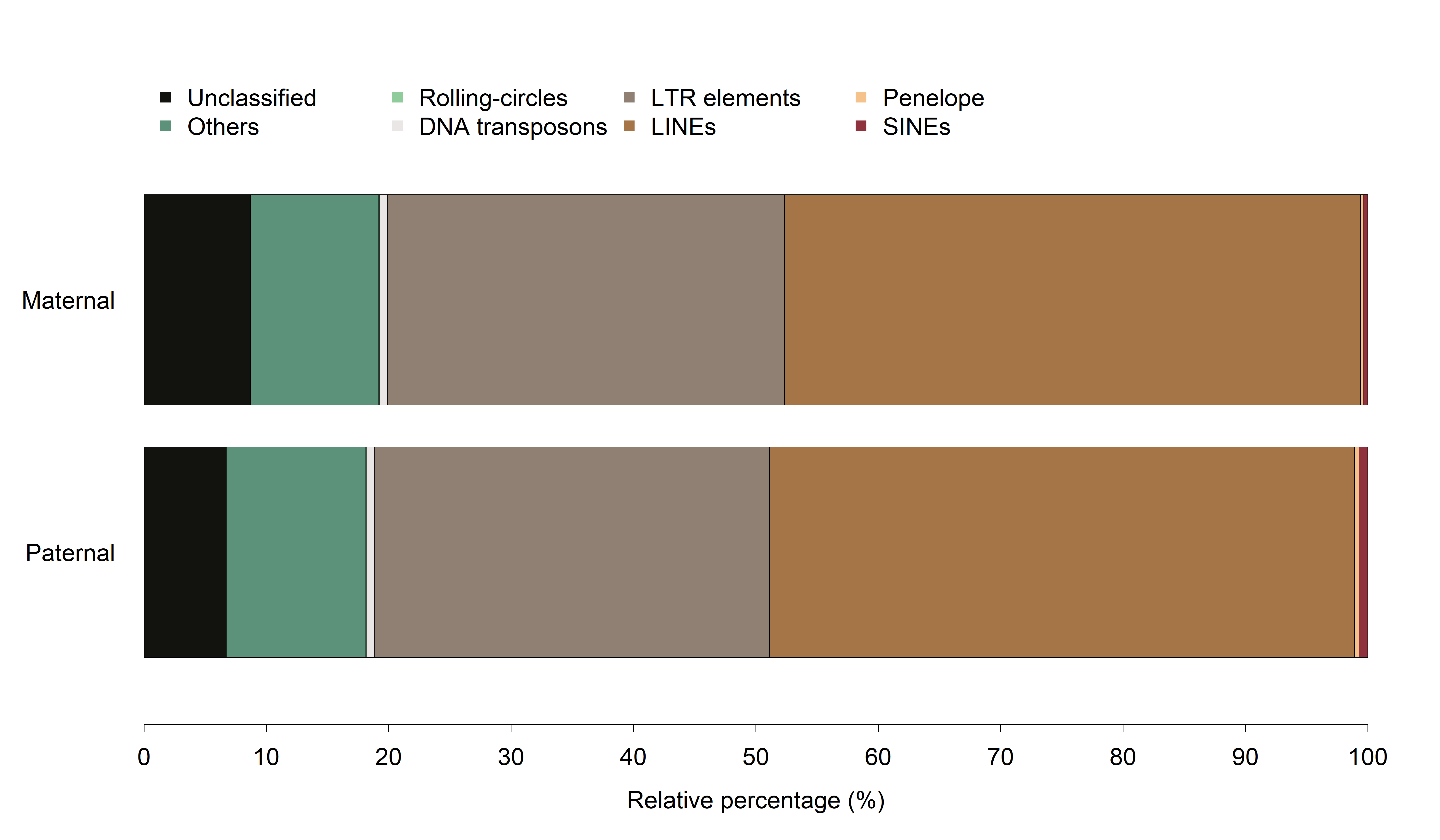

Figure S3

**Repeat landscape of the paternal and maternal haplomes.** Proportions of repeat classes annotated with RepeatModeler and classified with RepeatMasker are shown for each parental haplome. Percentages are expressed relative to the entire repeat content of the corresponding haplome.

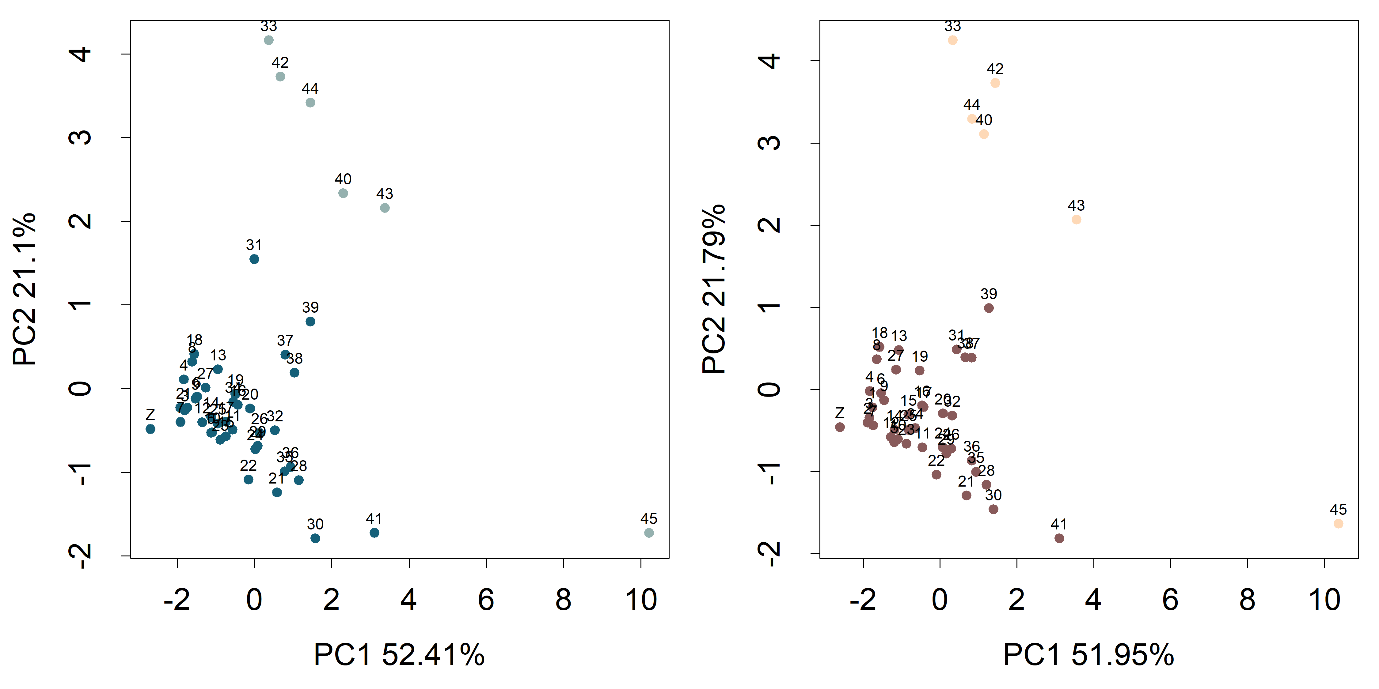

Figure S4

**PCA of chromosome level features (z-scored).** Principal component analysis was performed on the following features: (i) length, (ii) total exonic length (relative), (iii) GC content, (iv) gene density, (v) exon density, (vi) mean exon length, (vii) fraction of simple/low complexity repeats, and (viii) fraction of other RepeatMasker classes (overlaps resolved by the larger element’s class). Paternal chromosomes are shown in the left panel and maternal chromosomes in the right panel; chromosomes absent from the previous reference (GCF_018691265.1) are highlighted in lighter colours in both panels.

### Supplementary Tables

| **File name** | **Read count** | **Average read length (bp)** |
| --- | --- | --- |
| m64046_200313_184523.hifi_reads.fa.gz | 1633571 | 14023 |
| m64046_200512_231235.hifi_reads.fa.gz | 1884925 | 13882 |
| m64156_220708_235329.hifi_reads.fa.gz | 1636736 | 23046 |
| m64156_220712_124431.hifi_reads.fa.gz | 1639724 | 22774 |
| m64156_230805_122943.hifi_reads.fa.gz | 1289514 | 20680 |
| ON1_25k.fq.gz | 1500988 | 48702 |
| ON2_25k.fq.gz | 755044 | 52234 |

Table S1

**Summary of sequencing libraries.** PacBio HiFi and ONT libraries received from the Lausanne Genomic Technologies Facility (GTF) and the Gene Expression Core Facility (GECF), respectively, with the corresponding read count and mean read length (bp) for each library. In this table, the five HiFi libraries are listed first, followed by the two ONT libraries.

Table S2

**Chromosome-level features of both haplomes.** Are listed (i) length, (ii) total exonic length (relative), (iii) GC, (iv) gene density, (v) exon density, (vi) mean exon length; (vii) fraction of simple/low complexity repeats and (viii) fraction of other RepeatMasker classes (resolving overlaps by the larger element’s class).

| **Chr** | **Paternal** | | | | **Maternal** | | | |
| --- | --- | --- | --- | --- | --- | --- | --- | --- |
|  | **Telomere** | **Length** | **Pericentromeres** | **Length** | **Telomere** | **Length** | **Pericentromeres** | **Length** |
| 1 |  |  | X | 23 | X | 9851 |  |  |
| 2 |  |  |  |  | X | 604 |  |  |
| 3 |  |  | X | 2132 | X | 5956 | X | 43 |
| 4 |  |  | X | 133 | X | 8967 |  |  |
| 5 | X | 4041 |  |  |  |  | X | 9851 |
| 6 | X | 8905 |  |  | X | 9958 |  |  |
| 7 |  |  |  |  | X | 9042 |  |  |
| 8 | X | 9797 |  |  | X | 9871 |  |  |
| 9 | X | 1712 |  |  |  |  | X | 329 |
| 10 |  |  | X | 358 |  |  |  |  |
| 11 | X | 9941 |  |  | X | 9970 |  |  |
| 12 |  |  |  |  | X | 305 |  |  |
| 13 | X | 3502 |  |  | X | 7406 |  |  |
| 14 | X | 9728 |  |  |  |  | X | 30 |
| 15 | X | 7262 |  |  |  |  |  |  |
| 16 | X | 1234 | X | 13 |  |  | X | 342 |
| 17 |  |  | X | 52 | X | 6892 | X | 23 |
| 18 | X | 9670 | X | 25 | X | 6243 |  |  |
| 19 | X | 9656 |  |  | X | 5730 |  |  |
| 20 |  |  | X | 153 | X | 8895 | X | 160 |
| 21 | X | 7829 | X | 401 |  |  | X | 128 |
| 22 |  |  | X | 4594 | X | 7332 | X | 361 |
| 23 | X | 8689 |  |  | X | 2285 | X | 25 |
| 24 | X | 7844 | X | 9507 | X | 8253 | X | 167 |
| 25 | X | 8348 | X | 357 |  |  | X | 2598 |
| 26 | X | 9287 | X | 116 | X | 8362 | X | 318 |
| 27 | X | 172 |  |  | X | 9201 |  |  |
| 28 | X | 2974 | X | 280 | X | 4290 | X | 75 |
| 29 | X | 5996 | X | 3163 |  |  | X | 2513 |
| 30 | X | 9934 |  |  | X | 7646 |  |  |
| 31 | X | 6758 |  |  | X | 6979 |  |  |
| 32 | X | 1422 | X | 3592 | X | 8103 |  |  |
| 33 | X | 8004 | X | 16764 | X | 7837 |  |  |
| 34 | X | 4780 | X | 4938 | X | 9683 | X | 6450 |
| 35 |  |  | X | 4077 | X | 9814 |  |  |
| 36 | X | 1686 |  |  | X | 9191 |  |  |
| 37 | X | 8261 |  |  | X | 9062 |  |  |
| 38 | X | 6664 |  |  | X | 1080 |  |  |
| 39 | X | 7856 |  |  | X | 824 |  |  |
| 40 |  |  |  |  |  |  |  |  |
| 41 | X | 5455 |  |  | X | 2237 |  |  |
| 42 |  |  |  |  |  |  | X | 344 |
| 43 | X | 9947 | X | 244 | X | 754 | X | 93 |
| 44 |  |  |  |  |  |  |  |  |
| 45 | X | 2784 | X | 9570 | X | 3381 |  |  |
| 46 |  |  |  |  | X | 5252 |  |  |

Table S3

**Terminal features per chromosome.** For each parental chromosome, presence/absence of telomeric and pericentromeric sequence and the corresponding span (bp).

| **Set of 49-mers** | **Distinct** | **Unique** | **Total** | **Maximum count** |
| --- | --- | --- | --- | --- |
| father.jf49.L1.Uinf.selection1 | 3594562817 | 2096246897 | 9068873521 | 9273511 |
| father.jf49.L1.Uinf.selection2 | 3594043171 | 2095620581 | 9069098779 | 9290946 |
| father.jf49.L1.Uinf.selection3 | 3561856628 | 2066495849 | 9060796066 | 9632777 |
| mother.jf49.L1.Uinf.selection1 | 3116725701 | 1568256693 | 9079234142 | 5416817 |
| mother.jf49.L1.Uinf.selection2 | 3116301751 | 1567585954 | 9079153908 | 4382415 |

Table S4

**Counts and composition of parental 49-mer libraries.** For each parental 49‑mer library (three paternal, two maternal), we report the number of distinct 49‑mers, the number of unique 49‑mers, the total 49‑mer observations, and the maximum per‑49‑mer copy count.

| **Parent specific 49-mers set** | **Distinct** | **Total** | **Maximum count** |
| --- | --- | --- | --- |
| only_father.selection1.NOTmother1and2 | 179277071 | 548680948 | 3026 |
| only_father.selection2.NOTmother1and2 | 179332652 | 548753287 | 3086 |
| only_father.selection3.NOTmother1and2 | 180935695 | 556322694 | 3132 |
| only_mother.selection1.NOTfather1and2 | 188431608 | 622399767 | 728 |
| only_mother.selection1.NOTfather1and3 | 188853699 | 623561440 | 513 |
| only_mother.selection1.NOTfather2and3 | 188821563 | 623729275 | 613 |
| only_mother.selection2.NOTfather1and2 | 188549060 | 622942205 | 797 |
| only_mother.selection2.NOTfather1and3 | 188929355 | 624037708 | 483 |
| only_mother.selection2.NOTfather2and3 | 188912686 | 624182568 | 511 |

Table S5

**Parent-private 49-mers.** Paternal and maternal 49mer sets used identify parent-specific sequences. 49mers contain no Ns and are considered parent-private when present ≥ 2× in a subset of one parent and absent from subsets of the other parent.

| **Categories** | **Father** | **Mother** | **Both/Unclassified** |
| --- | --- | --- | --- |
| Centromere-like | 12,575 | 7,450 | 5,411 |
| Kinetochore-like | 187,463 | 37,388 | 8,132 |
| Microsatellite | 116,684 | 117,378 | 16,532 |
| Telomere | 3,273 | 2,876 | 125 |
| Other | 3,714,919 | 3,626,898 | 227,366 |

Table S6

**HiFi reads parental assignment by category.** The number of paternal, maternal and unclassified reads for each of the read category determined by their repeat profile (see *Identification of repeated sequences and read filtering* section for details).

| **Categories** | **Father** | **Mother** | **Both/Unclassified** |
| --- | --- | --- | --- |
| Centromere-like | 4,635 | 2,069 | 1,366 |
| Kinetochore-like | 74,039 | 17,016 | 5,305 |
| Microsatellite | 18,940 | 20,744 | 2,405 |
| Telomere | 1,173 | 1,040 | 98 |
| Other | 438,778 | 437,726 | 103,389 |

Table S7

**ONT (>= 25k) reads parental assignment by category.** The number of paternal, maternal and unclassified reads for each of the read category determined by their repeat profile (see *Identification of repeated sequences and read filtering* section for details).
